# Bayesian adaptive stimulus selection for dissociating models of psychophysical data

**DOI:** 10.1101/220590

**Authors:** James R. H. Cooke, Luc P. J. Selen, Robert J. van Beers, W. Pieter Medendorp

## Abstract

Comparing models facilitates testing different hypotheses regarding the computational basis of perception and action. Effective model comparison requires stimuli for which models make different predictions. Typically, experiments use a predetermined set of stimuli or sample stimuli randomly. Both methods have limitations; a predetermined set may not contain stimuli that dissociate the models whereas random sampling may be inefficient. To overcome these limitations, we expanded the psi-algorithm (Kontsevich & Tyler, 1999) from estimating the parameters of a psychometric curve to distinguishing models. To test our algorithm, we applied it to two distinct problems. First, we investigated dissociating sensory noise models. We simulated ideal observers with different noise models performing a 2-afc task. Stimuli were selected randomly or using our algorithm. We found using our algorithm improved the accuracy of model comparison. We also validated the algorithm in subjects by inferring which noise model underlies speed perception. Our algorithm converged quickly to the model previously proposed (Stocker & Simoncelli, 2006), whereas if stimuli were selected randomly model probabilities separated slower and sometimes supported alternative models. Second, we applied our algorithm to a different problem; comparing models of target selection under body acceleration. Previous work found target choice preference is modulated by whole body acceleration (Rincon-Gonzalez et al., 2016). However, the effect is subtle making model comparison difficult. We show that selecting stimuli adaptively could have led to stronger conclusions in model comparison. We conclude that our technique is more efficient and more reliable than current methods of stimulus selection for dissociating models.

**Data Availability:** All data and code will be posted on our institutional repository system following acceptance. In the meantime feel free to contact the authors if you would like any of the code.

## Introduction

Within neuroscience there is a clear interest in developing computational models to explain neural systems and behavior. This is seen in many disciplines, such as working memory (Keshvari, van den Berg, & Ma, 2012, 2013), speed perception (Stocker & Simoncelli, 2006), multisensory integration (Acerbi, Dokka, Angelaki, & Ma, 2017; Kording et al., 2007), effector selection (Bakker, Weijer, van Beers, Selen, & Medendorp, 2017), contrast gain tuning (DiMattina, 2016), and temporal interval reproduction (Acerbi, Wolpert, & Vijayakumar, 2012).

Inferring the best model out of several proposed models is important. Unfortunately, model comparison is typically difficult. In addition to the computational problem of having to integrate over the parameter space of each model, it is also necessary to present stimuli which can dissociate the models. If different psychophysical models make similar predictions for many of the stimuli presented then it is difficult to dissociate these models. Despite the importance of appropriate stimuli selection many studies comparing models either select stimuli randomly (Keshvari et al., 2012, 2013) or use a set of constant stimuli (Acerbi et al., 2012, 2017; Bakker et al., 2017; Kording et al., 2007). Both of these approaches may select stimuli that are uninformative for model comparison, resulting in a large number of trials to accurately distinguish different models.

A more efficient approach is to select stimuli that optimize some criterion (often referred to as a utility function). The idea of utility-based stimulus selection has been studied extensively in statistics and machine learning, typically called active learning (Gardner et al., 2015; Kulick, Lieck, & Toussaint, 2014), adaptive design optimization (Cavagnaro, Myung, Pitt, & Kujala, 2010) and optimal experiment design (DiMattina & Zhang, 2011). These types of algorithms have been applied to a wide range of problems including neuronal tuning curve estimation (Pillow & Park, 2016), testing for deficits in auditory perception (Gardner et al., 2015), and machine classification (Houlsby, Husz´ar, Ghahramani, & Lengyel, 2011) but are not commonly employed in psychophysics. For a more comprehensive review on the application of adaptive stimulus selection in sensory systems neuroscience see DiMattina and Zhang (2013).

Within psychophysics, selecting stimuli in an adaptive manner has been used extensively for estimating the parameters of a specific psychophysical model. For example, Kontsevich and Tyler (1999) used an information theoretic approach to estimate the slope and threshold parameters of a one-dimensional psychometric function, selecting on each trial the stimulus which maximizes the information gain about these parameters. Additional work then improved on this by marginalizing out unwanted parameters in order to improve the estimates of desired parameters (Prins, 2013). However, many psychophysical models are not uni-dimensional and as such this approach was extended to multi-dimensional models (DiMattina, 2015; Kujala & Lukka, 2006; Lesmes, Lu, Baek, & Albright, 2010).

What if instead of inferring the parameters of these multidimensional models, we wish to dissociate different models? Wang and Simoncelli (2008) developed an algorithm specifically designed for generating stimuli on a trial-to-trial basis to compare two psychophysical models. However, in many cases there are more than two candidate models. More recent work used an information theoretic approach to derive a method for optimal stimulus selection to compare an arbitrary number of models (DiMattina, 2016). However, this approach does not determine the optimal stimulus on a trial-to-trial basis and therefore may be a suboptimal approach. Recently, a general approach for determining the optimal stimulus to compare multiple models has been proposed in the field of cognitive science (Cavagnaro et al., 2010; Cavagnaro, Pitt, & Myung, 2011; Cavagnaro, Gonzalez, Myung, & Pitt, 2013). This approach, named Adaptive Design Optimization (ADO), which simulates the utility distribution of possible stimuli, can be done on a trial-to-trial basis (Cavagnaro et al., 2013) and could be used to distinguish more than two models. This makes it a potentially powerful tool to select stimuli for comparing models of psychophysical data. However, implementing this approach requires a detailed understanding of Monte Carlo based simulation approaches such as particle filtering and simulated annealing.

This difficulty may prohibit widespread adoption of ADO. Therefore, we present an alternative and easier to implement algorithm for selecting stimuli on a trial-to-trial basis to dissociate multiple models of psychophysical data. The algorithm is a generalization of the classical psi-method (Kontsevich & Tyler, 1999; Prins, 2013), shifting from estimating parameters of models to comparing models. In order to test our algorithm, we applied it to two very different psychophysical problems. First, we tested dissociating distinct models of sensory noise which affect speed perception. In order to do this we constructed three generative models, each with its own noise properties, that were probed by an ideal observer performing a 2-afc task. Stimuli were either selected randomly, using our adaptive algorithm or using a more classical approach of measuring psychometric curves around a variety of fixed references. We found that when stimuli are selected adaptively, the accuracy of model comparison improved. We also tested our algorithm in real subjects by inferring which of three sensory noise models best explains their behavior in a speed perception task. To do this, we used a psychophysical experiment in which stimuli were either selected randomly or adaptively. The adaptive procedure converged to the model proposed in earlier work (Stocker & Simoncelli, 2006) whereas the random sampling method was often inconclusive about the underlying noise model. Second, we tested the algorithm on dissociating two models of saccadic target selection under whole body acceleration (Rincon-Gonzalez et al., 2016). Based on the original experimental data it is hard to dissociate between an acceleration-dependent or acceleration-independent target selection model at the individual subject level. However, using simulations, we show that selecting the stimuli adaptively could have led to stronger conclusions during model comparison. We conclude that our technique is more accurate and faster than the current methods to dissociate psychophysical models. In addition, we provide a python implementation of our algorithm, as well as the code and data to perform the simulations and analysis presented.

## Methods

Our algorithm is based on an experimenter wishing to determine which of a set of *m* discrete psychophysical models best describes subject’s behavior, under the assumption that the model underlying subjects behavior is contained in the set of models. Under a traditional experimental approach an experimenter would present a number of stimuli **x** to a subject and obtain the corresponding responses to these stimuli **r**. Using Bayes’ rule, we can compute the probability of a particular psychophysical model *m* given the responses and stimuli as:

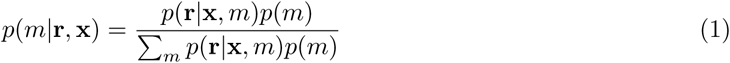

where *p*(*m*) is the prior probability of each model *m*, *p*(*m*|**r**, **x**) is the posterior distribution of each model and *p*(**r**|**x**,*m*) is referred to as the marginal likelihood. The marginal likelihood is obtained by marginalizing over the parameters *θ* of the particular model:

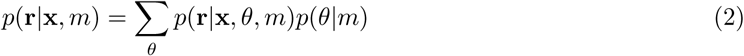

Equation 1 makes it clear that our ability to dissociate models is dependent on the stimuli **x** that were presented to the subject. Different stimuli and responses produce different posterior distributions of models. We can characterize the quality of a possible posterior using a particular utility function. Following previous work in model comparison, we use the entropy of the posterior distribution to characterize its quality (Cavagnaro et al., 2010, 2011, 2013; DiMattina, 2016):

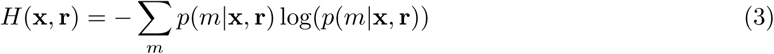

A posterior with lower entropy entails more certainty about which model underlies the subjects’ behavior. A minimal entropy distribution across models would be a posterior mass of 1 at a single model and 0 at all others.

How should we select stimuli to minimize the expected entropy of the model posterior? Here we propose using a similar approach to that used previously for minimizing the entropy of a parameter posterior (Kontsevich & Tyler, 1999), by numerically calculating on each trial the stimulus that minimizes the expected entropy of the model posterior. For our algorithm, we represent the possible stimuli on each trial *x* and parameters *θ* on discrete grids, similar to Kontsevich and Tyler (1999). This requires three quantities: a prior distribution over models *p*(*m*), a prior distribution of parameters for each model *p*(*θ*|*m*), and a likelihood look-up table for each model *p*(*r*|*x, θ, m*) which represents the probability of a response given a model and parameter set. Using these quantities, we can design an iterative algorithm to select the optimal stimuli on a trial-to-trial basis, which is as follows:

1. Calculate for each model and all possible stimuli the marginal likelihood of a response at trial *t* given stimulus *x*:

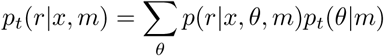
2. Compute the posterior distribution of models given response *r* in the next trial to stimulus *x*:

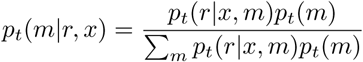 Note, Σ_*m*_ *p_t_*(*r*|*x, m*)*p_t_*(*m*) can also be written *p_t_*(*r*|*x*) and should be stored as the term is also used in step 4.
3. Compute the entropy of the posterior distribution over models given presented stimulus *x* and response *r*:

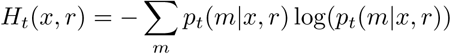
4. Because the response is unknown before the trial, we must marginalize over all possible responses to obtain the expected entropy:

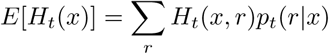
5. Find the stimulus that produces a posterior with the minimum expected entropy:

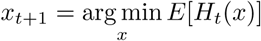
6. Use *x*_*t*+1_ as the stimulus on the next trial to receive response *r*_*t*+1_.
7. Because step 1 requires a prior on the parameters *p*_*t*_(*θ*|*m*), this prior must be recursively updated in addition to updating the model priors. As such we set the parameter and model priors to their posteriors:

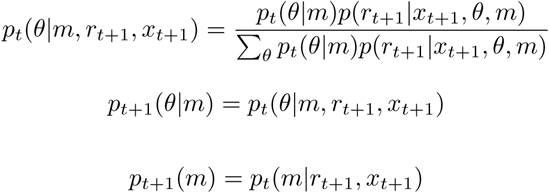
8. Return to the first step until the desired number of trials is completed or sufficent model evidence has been obtained.

## Experiment 1: Velocity judgment

### Introduction

Most computational models of perception and action take one particular assumption about how the sensory uncertainty depends on the stimuli presented. For example, there are models that assume sensory noise is constant and independent of the stimuli presented (Kording et al., 2007; Weiss, Simoncelli, & Adelson, 2002), some assume a linear increase in the standard deviation of the noise with the stimulus magnitude (Battaglia, Kersten, & Schrater, 2011; Sanborn & Beierholm, 2016), others take a combination of these two (Odegaard, Wozny, & Shams, 2015; Petzschner & Glasauer, 2011; Stocker & Simoncelli, 2006). To our knowledge only a few papers made an explicit comparison between sensory noise models (Acerbi et al., 2012, 2017; Jazayeri & Shadlen, 2010). A striking finding in these comparison studies is that the sensory noise model can vary among subjects (Acerbi et al., 2012, 2017). Given that the predictions of complex models, for example, models of multisensory integration (Acerbi et al., 2017) are dependent on the assumed sensory noise model, it is important to have an accurate model of each subject’s sensory noise model. It is therefore essential to validate the assumed sensory noise model to ensure it is accurate.

One way to validate these assumptions is by performing an additional experiment designed to estimate the observer’s sensory noise model. However, performing an additional experiment requires more time and resources. Being able to minimize the number of trials required to perform this type of comparison (as well as increasing the inference accuracy) is therefore beneficial. This presents a potential use of our algorithm, a method to validate sensory noise models and infer them for use in more complex models. Here, we use both simulation and a behavioral experiment to demonstrate that our algorithm can be used to facilitate inference of a subject’s sensory noise model. More specifically, as an illustrative example, we focus on inferring the sensory noise model underlying speed perception. We used this paradigm for two reasons. First, it is experimentally quick to test so we can compare our algorithm to other methods of stimuli selection. Second, previous work assumed a sensory noise model which consisted of both a constant component (the sensor is not perfect even when speed is zero) and a component that linearly increases with speed (Stocker & Simoncelli, 2006) and thus we can compare our inference to this model.

## Methods

### Models

In order to test between different sensory noise models we need to specify a model of the subjects’ responses. We derived a simple 2-afc model of subject responses using signal detection theory (see appendix A). This leads to the response probability given a probe *s*_2_ and a reference *s*_1_, described by:

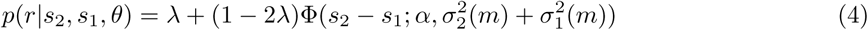

in which Φ is the cumulative density function of a Gaussian distribution, evaluated at point *s*_2_ − *s*_1_ with a mean *α* and variance 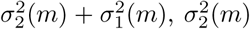 and 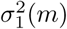 are the variances of the sensory noise for the probe and reference stimuli respectively, *λ* is a lapse rate accounting for trials where an observer guesses randomly, and *α* is a bias parameter accounting for biases in subject’s responses. We assume the subject’s sensory noise changes with the stimulus in one of three ways. The first, and simplest model, assumes sensory variance is independent of the stimulus. We denote this the constant noise model. The second model assumes that the standard deviation of the sensory noise increases linearly with the signal intensity, and thus has zero standard deviation if the signal is absent. This model is referred to as the Weber model. Finally, we consider a model where the sensory noise is non-zero when the signal is absent and also has a linearly increasing part, which we will refer to as the generalized model.

For the constant model, we assume the sensory variance is constant *σ*^2^ = (5*β*)^2^ (this parameterization allows *β* to be kept in a similar range for each model), for the Weber model we assume *σ*^2^ = (*βs*)^2^, and for the generalized model we assume *σ*^2^ = *γ*^2^ + (*βs*)^2^. The above response model means we can parametrize a subject’s response behavior (regardless of model) using 4 parameters, *θ* = [*α, β, γ, λ*].

### Simulation experiment

In order to investigate whether using our adaptive algorithm facilitates comparison of sensory noise models, we first performed a simulation experiment. To this end, we need to specify the grids to use for the stimuli and parameters as well as the priors. The lower bound, upper bound, and number of steps for all variables are shown in Table 1. For the prior over parameters *p*(*θ*|*m*), we assumed a uniform discrete distribution for each parameter and that the parameters are independent. Finally, for the prior over models *p*(*m*) we used a uniform distribution over the three models.

**Table 1.** Parameter grids used for simulation experiment 1 and the adaptive and random conditions in our subject experiment. *s*_1_ is the reference speed stimulus, *s*_2_ is the probe speed stimulus, *α* is a bias parameter, *β* is a scaling parameter for the subject’s sensory uncertainty, *γ* is the base sensory uncertainty of an observer (only used in the generalized model), and *λ* is the lapse rate of an observer.

| Variable | Lower bound grid | Upper bound grid | Number of steps |
| --- | --- | --- | --- |
| $s_1$ (deg/s) | 0.6 | 9 | 10 |
| $s_2$ (deg/s) | 0.3 | 9 | 20 |
| $\alpha$ (deg/s) | -0.6 | 0.6 | 17 |
| $\beta$ (-) | 0.01 | 0.5 | 25 |
| $\gamma$ (deg/s) | 0 | 2 | 20 |
| $\lambda$ (-) | 0 | 0.1 | 10 |

As different subjects could have different parameters and noise models it is important to test our algorithm over a wide range of parameters and models. As such, we first generated 2000 possible parameter combinations. The parameters were drawn independently from a continuous uniform distribution with the same upper and lower bounds as those specified in Table 1. Next, in order to assess how well we can infer the correct generative model, we simulated 750 trials from each model for each parameter combination. This entailed using the same parameter combination for each model (as the constant and Weber models are not dependent on *γ*, it was not used for these models). The stimuli for these trials were either selected adaptively using our algorithm, or randomly from the same stimulus grid. This led to a total of 12000 simulated datasets.

We used uniform priors to match the uniform distribution we drew our parameters from. In practice any prior distribution could be used, but if it is continuous, the grid representation will create a discrete approximation. We also performed an additional simulation using a truncated Gaussian parameter distribution (Supplemental material) to better asses the performance of our algorithm.

### Real experiment

We also tested whether our algorithm could facilitate model comparison in actual subjects. This was done using a 2-afc speed judgment task in which stimuli were selected in one of three ways, adaptively (using our algorithm), randomly (from the same stimulus grid as adaptive), or using the traditional approach of measuring separate psychometric curves for different reference values (Stocker & Simoncelli, 2004, 2006) using the psi algorithm (Kontsevich & Tyler, 1999). We tested 6 naive subjects (4 female, aged 25-34). The experiment was approved by the local ethics committee of the Social Sciences Faculty of Radboud University. In accordance with the Declaration of Helsinki written informed consent was obtained from all subjects prior to the experiment.

The stimuli consisted of two drifting Gabor patches and a black fixation dot, which were drawn using PsychoPy (Peirce, 2009). Both patches were 3 deg of visual angle in size, with a spatial frequency of 1.5 cycle/deg, the contrast of each was set to 90%, and the stimuli were drawn at 6 deg on either side of fixation. The background was grey with a luminance of 91.17 cd/m^2^. The fixation dot was 0.2 deg in size and drawn in the center of the screen. The stimuli were displayed at a resolution of 1024 by 768 on a gamma corrected 17 inch Iiyama HM903DTB monitor viewed from a distance of approximately 43.5 cm.

On each trial, the subject saw both Gabors drift simultaneously and horizontally for 1 s. Both Gabors moved in the same direction on a given trial (direction was left or right and was selected randomly for each trial). One Gabor (the reference) drifted with speed *s*_1_ deg/s and the other (the probe) with speed *s*_2_ deg/s. The subject was asked to judge which of the two was faster and indicate this with a button press. The position of the reference stimulus (left or right of fixation) was randomized on each trial. The experiment was split into two sessions, the ordering of which was counterbalances across subjects. In one session (algorithm session) subjects performed 1500 trials, 750 of which were adaptive trials and 750 were random trials. On an adaptive trial, the Gabor speeds were selected using our algorithm based on the previous stimuli (and responses) generated by this algorithm; on a random trial the speed of each Gabor was selected randomly from the stimulus grid. The stimuli and parameter grids used were the same as for the simulation experiment. In this session the screen was refreshed at 72 Hz.

In another session (psi session), subjects performed 750 trials designed to measure their psychometric curve for five reference values (150 trials per reference, see Table 2 for the reference values used). On each trial, *s*_1_ was randomly selected from a set of 5 possible values, the value of *s*_2_ on this trial was then selected using the psi-marginal algorithm (Prins, 2013) (see Table 2 for the grids used). This was done in order to maximize the information gain about *µ* (the point of subjective equality) and *σ* (the standard deviation) for this particular value of *s*_1_ under the assumption the probability of a subjects response follows:

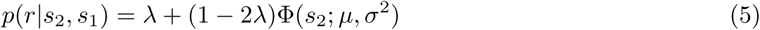

in this equation *σ*^2^ is the variance of the normal distribution and *µ* is the mean of the distribution. Selecting stimuli in this manner, allows us to assess how effective the more traditional fixed reference approach is to separating sensory noise models compared to our algorithm. In this session, stimuli were refreshed at 144 Hz. Note that the probe *s*_2_ had a denser grid in this session (see Table 2 compared to Table 1); this allows us to better estimate the psychometric curve of each subject but may also give an advantage to this method in terms of model comparison. Prior to each session, subjects performed 20 practice trials from the respective session.

**Table 2.** Parameter grids used in our fixed reference condition. *N*(*a, b*) indicates the prior was normally distributed with mean *a* and standard deviation *σ*, *U* indicates a discrete uniform distribution and *B*(*a, b*) indicates a beta distribution with shape parameters *a* and *b*. The values for *s*_1_ were determined based on previous work on speed perception (Stocker & Simoncelli, 2004). The prior for *µ* was selected based on the assumption that the psychometric curve for a 2-afc task will be close to unbiased. The prior for *λ* was selected based on recommendations for the psignifit toolbox (Fründ, Haenel, & Wichmann, 2011) (see http://psignifit.sourceforge.net).

| Variable | Lower bound grid | Upper bound grid | Number of steps | Prior | $s_1$ (deg/s) |
| --- | --- | --- | --- | --- | --- |
| $\mu$ (deg/s) | 0.001 | 3 | 41 | $N(0.5, 2)$ | 0.5 |
| $\sigma$ (deg/s) | 0.01 | 3 | 51 | $U$ | 0.5 |
| $\lambda$ (-) | 0 | 0.1 | 15 | $B(2, 20)$ | 0.5 |
| $s_2$ (deg/s) | 0.01 | 3 | 61 | N/A | 0.5 |
| $\mu$ (deg/s) | 0.001 | 4 | 41 | $N(1, 2)$ | 1 |
| $\sigma$ (deg/s) | 0.01 | 4 | 51 | $U$ | 1 |
| $\lambda$ (-) | 0 | 0.1 | 15 | $B(2, 20)$ | 1 |
| $s_2$ (deg/s) | 0.01 | 4 | 61 | N/A | 1 |
| $\mu$ (deg/s) | 0.1 | 6 | 41 | $N(2, 2)$ | 2 |
| $\sigma$ (deg/s) | 0.01 | 6 | 51 | $U$ | 2 |
| $\lambda$ (-) | 0 | 0.1 | 15 | $B(2, 20)$ | 2 |
| $s_2$ (deg/s) | 0.1 | 6 | 61 | N/A | 2 |
| $\mu$ (deg/s) | 1 | 9 | 41 | $N(4, 2)$ | 4 |
| $\sigma$ (deg/s) | 0.01 | 9 | 51 | $U$ | 4 |
| $\lambda$ (-) | 0 | 0.1 | 15 | $B(2, 20)$ | 4 |
| $s_2$ (deg/s) | 1 | 9 | 61 | N/A | 4 |
| $\mu$ (deg/s) | 0.001 | 14 | 41 | $N(4, 2)$ | 8 |
| $\sigma$ (deg/s) | 0.01 | 14 | 51 | $U$ | 8 |
| $\lambda$ (-) | 0 | 0.1 | 15 | $B(2, 20)$ | 8 |
| $s_2$ (deg/s) | 3 | 14 | 61 | N/A | 8 |

### Analysis

For our analysis, we used Python 2.7 (Python Software Foundation, https://www.python.org) and additional python based toolboxes, primarily SciPy (Jones, Oliphant, Peterson, & others, 2001), Numpy (Walt, Colbert, & Varoquaux, 2011), Matplotlib (Hunter, 2007), scikit-learn (Pedregosa et al., 2011) and Pandas (McKinney & others, 2010).

In addition to computing the model probabilities for every subject for the different sampling methods, we also estimated each subject’s parameters for each model by maximizing the log-likelihood of the parameter values based on the subject’s responses (to increase accuracy we pooled the data from all sessions). This provides more sensitive parameter estimates than the grid we used for model comparison and also allows us to check the parameters are not close to the edges of the grids we used.

We assumed the subject’s responses are independent across trials. The subject’s response probability on each trial can then be computed using equation 4. The log-likelihood of a parameter set given a subject’s entire data set, is given by

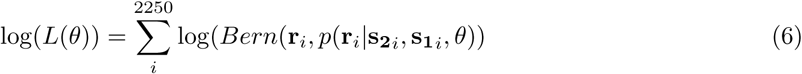

in which *i* is the trial index, **r** is a vector of subject response, **s_1_** is a vector of the reference stimuli, **s_2_** is a vector of probe stimuli and *Bern* stands for a Bernoulli distribution.

Parameter estimates 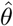 were then obtained by minimizing the negative log-likelihood:

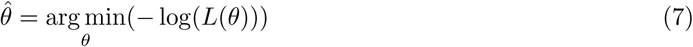

This optimization was done numerically using the L-BFGS-B algorithm (Byrd, Lu, Nocedal, & Zhu, 1995), implemented in SciPy (Jones et al., 2001) and applied in the *scipy.optimize.minimize* function. The L-BFGS-B is an iterative algorithm designed to optimize a nonlinear function subject to parameter boundaries (Byrd et al., 1995). The parameter bounds were set to those in Table 1. To ensure a global minimum was found we used 100 random initializations and selected the parameter set with the highest log-likelihood. The initial values were obtained by drawing each parameter value from a continuous uniform distribution with the same bound as those in Table 1.

In order to validate the results of the grid-based model comparison we also computed the Akaike Information Criterion (AIC) for each of the models. This is a metric which summaries how well a model fits (higher likelihood) the data while correcting for the number of parameters (Akaike, 1974; Burnham, Anderson, & Burnham, 2002),

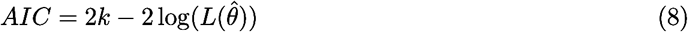

in which *k* is the number of parameters of the model. It is important to note that computing model probabilities using equation 1 also implicitly corrects for the number of parameters (MacKay, 2003).

## Results

### Simulation experiment

Figure 1 shows the model probabilities over trials averaged across the different parameter sets from our simulation experiment. As expected, the model probabilities trend towards 1 along the diagonal, indicating that both adaptive and random sampling converge towards the correct model. This demonstrates that our algorithm does not introduce any bias during model comparison, even when the number of parameters differs between models. It can also be observed that the probability of the correct model rises faster and is higher when we select stimuli adaptively (green curves) than when stimuli are selected randomly (orange curves). This indicates the strength of evidence towards the correct model is higher when we use adaptive sampling.

**Figure 1.**
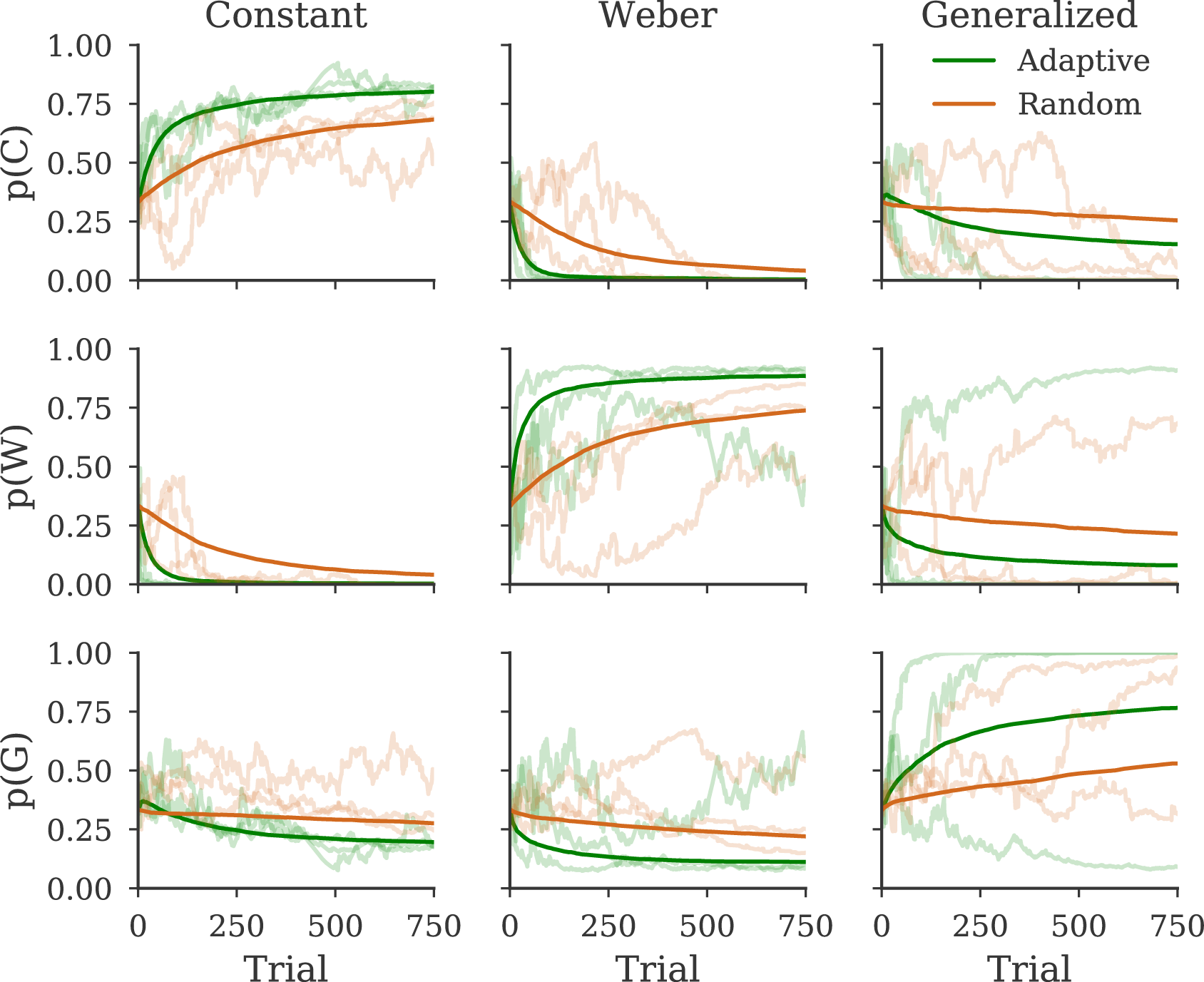
Evolution of model probabilities over trials for different generative models and algorithms. Columns indicate the model used to generate the data, rows indicate the probability of each model. The dark lines indicate the mean probability averaged over simulations, light lines indicate example simulations. Green coloring indicates stimuli were selected adaptively, orange coloring indicates stimuli were selected at random from the same stimulus grid.

Although Figure 1 provides evidence that adaptive sampling improves the strength of evidence towards the correct model, it does not quantify how this increase would affect the conclusions of an experiment. In order to quantify the practical benefit of adaptive sampling, we computed the ratio of the probability of the generative model against the other models (commonly referred to as the Bayes factor). This ratio represents how much more probable one model is than the other model (MacKay, 2003). Because we consider three models, this yields two Bayes factors, which the experimenter can use to decide whether there is significant evidence in favor of a particular model. A commonly used criterion is a that a Bayes factor over 3 indicates positive evidence towards this model (Kass & Raftery, 1995).

Figure 2 shows the proportion of simulations where the Bayes factors for the correct model against the other two models were both over 3. This represents the proportion of simulations in which we would find evidence in favor of the correct model. We see that adaptive sampling has a higher proportion than random sampling, indicating an experimenter would conclude in favor of the correct model more often using adaptive sampling. For example, an experimenter would be twice as likely to find strong evidence in favor of the correct model using our approach if the underlying model was the generalized one.

**Figure 2.**
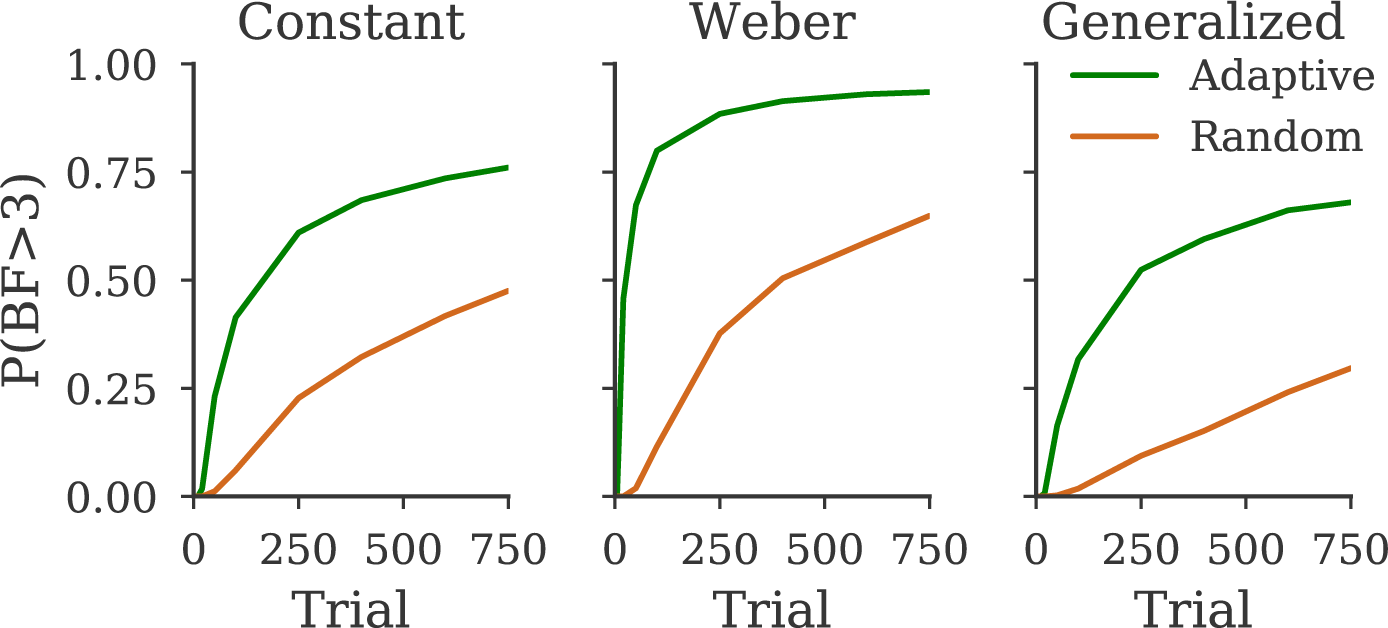
Proportion of simulations where both Bayes factors of the generative model relative to an alternative model is over 3, plotted as a function of the number of trials. Each column indicates the model used to generate the data.

While Figure 2 shows that adaptive sampling increases the probability of concluding in favor of the true generative model, it is not apparent why the proportion of Bayes factors over 3 is lower when stimuli are selected randomly. One possibility is that random sampling still supports the true generative model but the strength of this support is insufficient; another possibility is that random sampling supports the incorrect model.

In order to explore these possibilities we plotted the probability of the correct model for each sampling method as a function of *β* and *γ* (see Figure 3). Figure 3 shows that the model probabilities are primarily green to yellow when the generative model is Weber or Constant. This indicates both methods mostly select the correct model. We can also see that in general adaptive sampling produces model probabilities which trend closer to 1 (i.e. yellow) indicating stronger evidence in favor of the correct model. When the generative model is the Generalized model, a substantial number of simulations produce probabilities supporting alternative models (indicated by the blue shading). At first this seems counter intuitive. However, for small values of *γ* and *β*, the generalized model becomes almost equivalent to the Weber and constant model. Because these have fewer parameters, they are favored in this situation.

**Figure 3.**
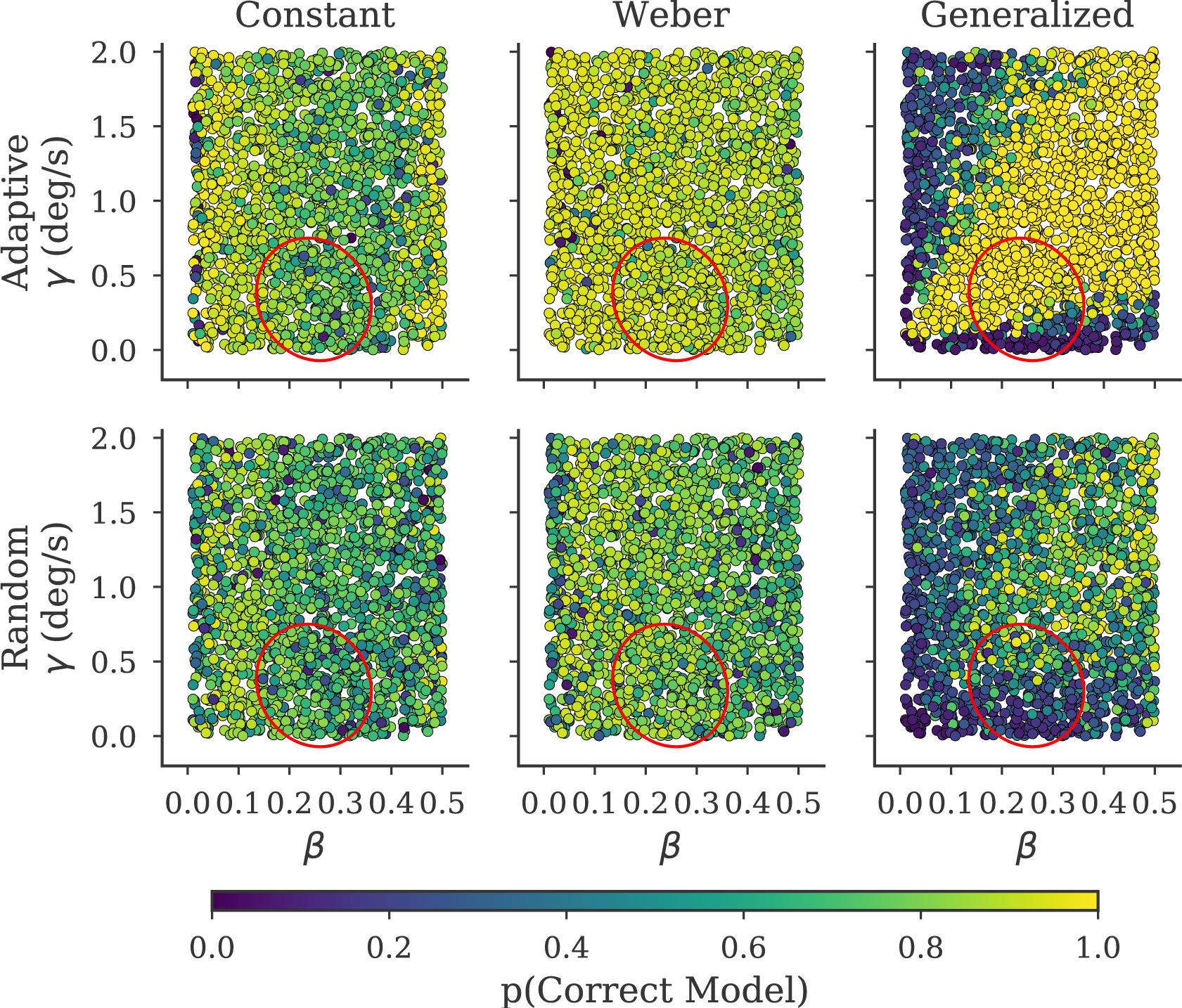
Probability of the generative model as a function of parameter values for different generative models and algorithms. Columns indicate the model used to generate the data, rows indicate the sampling method used to determine stimuli. Each point indicates the probability of the correct model as a function of the parameters *γ* and *β* for one simulation. Note, the Weber and constant models are independent of *γ* and thus model probabilities do not change systematically as a function of *γ*. The *γ* value plotted refers to the *γ* used in the generalized model for this simulation, all other parameters are shared between the models. The red ellipse indicates the mean ± two standard deviations of the subjects’ parameter estimates for *γ* and *β* obtained from the generalized model (see Table 3).

### Actual experiment

The previous section suggests that, in simulation, adaptive sampling provides a large benefit to model comparison. We next tested whether this improvement also transfers to actual experiments. Figure 4 shows the model probabilities of each subject obtained from our speed perception experiment and the average across subjects. As shown, on average adaptive sampling supports the generalized model, which is consistent with previous work (McKee, Silverman, & Nakayama, 1986; Stocker & Simoncelli, 2006). By contrast, both random sampling and sampling from the psi algorithm are indecisive as to the underlying noise model. The reason follows from inspecting the individual subject data. When stimuli are selected adaptively, the probability of the generalized model is high for all subjects. By contrast, random sampling supports the Weber model for 3 subjects and the generalized for the others (although the probability is lower than that found from adaptive sampling). The psi session provides similar results to the random session; 3 subjects are best described by a generalized model and the remaining by the Weber model. Given that the findings of the different sampling methods are disparate, we also computed AIC values on the data of all sessions grouped together, which allows us to assess which model is the best based on the entire data set (see Table 3). Shown by this table, the AIC results favor the generalized model for every subject, indicating that the results of the adaptive sampling method are comparable to the results of the grouped data. In addition, to asses the possibility that our adaptive technique was supporting the incorrect model, we performed additional simulations to verify that the observed differences between the sampling methods are as expected. Indeed, when the data is generated from the generalized model, the random sampling method often converges to the wrong, i.e Weber model (see Supplementary material). Together this suggests the conclusions drawn from the adaptive sampling method are more accurate than conclusion drawn from both random sampling or measuring independent psychometric curves.

**Table 3.** Best fit parameters and AIC (∆AIC, generalized - other model) of each model and subject.

| Subject | Model | $\alpha$ (deg/s) | $\beta$ (-) | $\lambda$ (-) | $\gamma$ (deg/s) | $\Delta AIC$ |
| --- | --- | --- | --- | --- | --- | --- |
| 1 | Weber | -0.043 | 0.354 | 0 | N/A | -17.459 |
| 1 | Constant | 0.075 | 0.073 | 0.1 | N/A | -202.656 |
| 1 | Generalized | 0.006 | 0.314 | 0 | 0.187 | 0 |
| 2 | Weber | -0.009 | 0.223 | 0.005 | N/A | -16.031 |
| 2 | Constant | 0.043 | 0.061 | 0.054 | N/A | -220.883 |
| 2 | Generalized | 0.016 | 0.197 | 0.004 | 0.122 | 0 |
| 3 | Weber | -0.136 | 0.268 | 0.055 | N/A | -171.633 |
| 3 | Constant | 0.027 | 0.195 | 0.027 | N/A | -141.811 |
| 3 | Generalized | 0.049 | 0.222 | 0.002 | 0.584 | 0 |
| 4 | Weber | -0.034 | 0.308 | 0.003 | N/A | -14.415 |
| 4 | Constant | 0.026 | 0.052 | 0.1 | N/A | -180.728 |
| 4 | Generalized | 0.002 | 0.272 | 0.004 | 0.161 | 0 |
| 5 | Weber | 0.019 | 0.256 | 0.023 | N/A | -109.731 |
| 5 | Constant | 0.074 | 0.182 | 0.007 | N/A | -151.102 |
| 5 | Generalized | 0.041 | 0.18 | 0 | 0.447 | 0 |
| 6 | Weber | -0.028 | 0.424 | 0 | N/A | -85.408 |
| 6 | Constant | 0.19 | 0.188 | 0.06 | N/A | -175.483 |
| 6 | Generalized | 0.149 | 0.303 | 0 | 0.533 | 0 |

**Figure 4.**
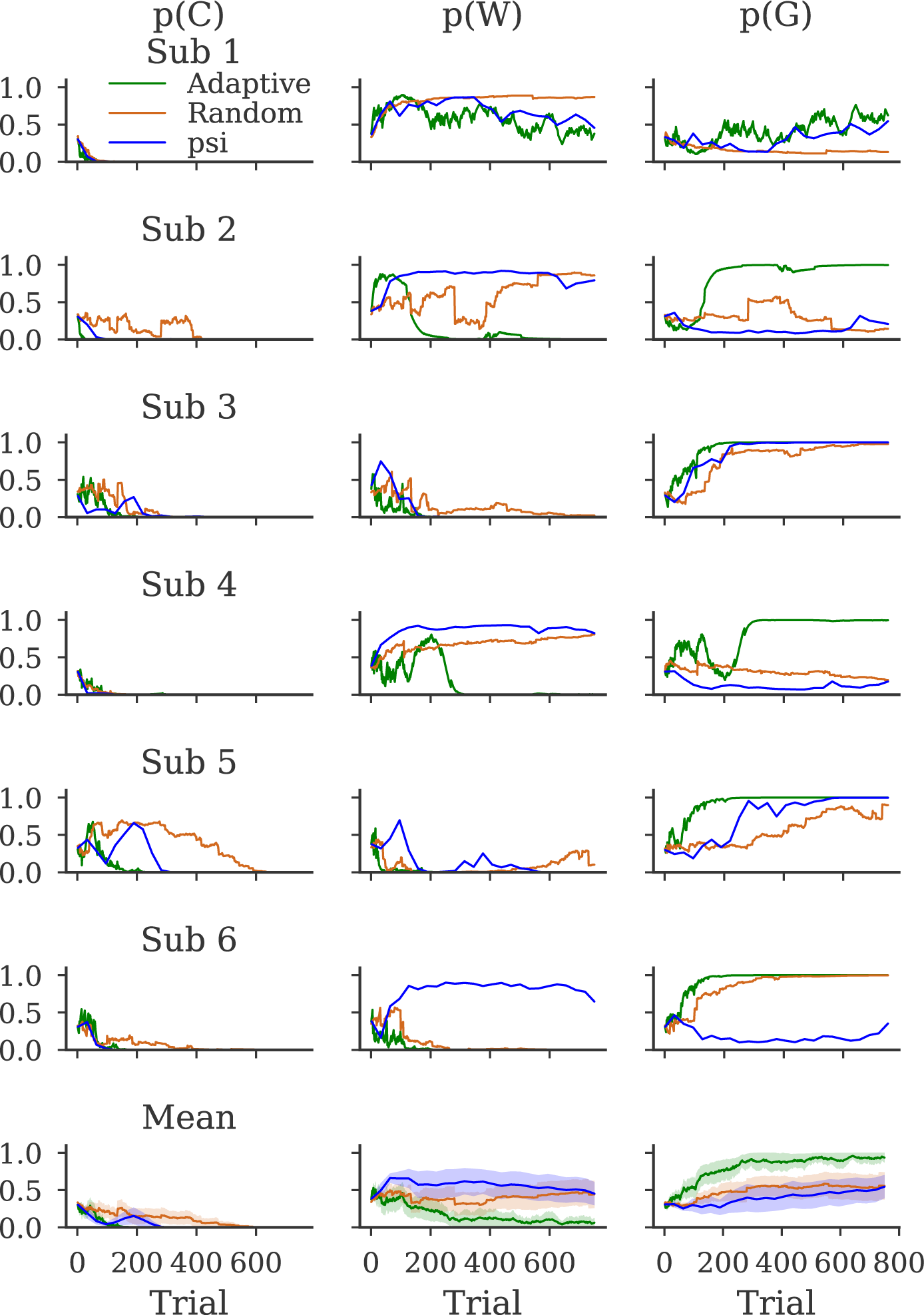
Evolution of model probabilities over trials for each subject. Columns indicate the probability of a particular model, rows indicate the subject. Green lines show the model probabilities when the stimuli were selected adaptively using our algorithm, orange lines indicate the model probabilities when stimulus were selected at random from the same stimuli grid, and blue lines indicate stimuli were selected using the psi algorithm. The lines in the mean plot show the mean model probabilities over subjects, the shaded area indicates *±* 1 *SEM* over subjects.

Although the results of the model comparison match previous work, it is important to note that a model being the most likely does not entail it fits the data well, just that it fits better than the other models. It is important to check the predictions of the models against the data.

Figure 5 illustrates the data of each subject obtained from the psi session as well as the predicted psychometric curves obtained from fitting the models to the data obtained from the adaptive algorithm only (therefore the models were not fit to the data shown). As shown, the constant model is in general a poor predictor of the data. By contrast, both the predictions of the Weber and generalized model are close to the data. This matches the results of AIC comparison (see Table 3) which indicated that the Weber and generalized model produce better fits to the data than the constant model. This also means that the assumptions with regards to our models (see Appendix A) are reasonable.

**Figure 5.**
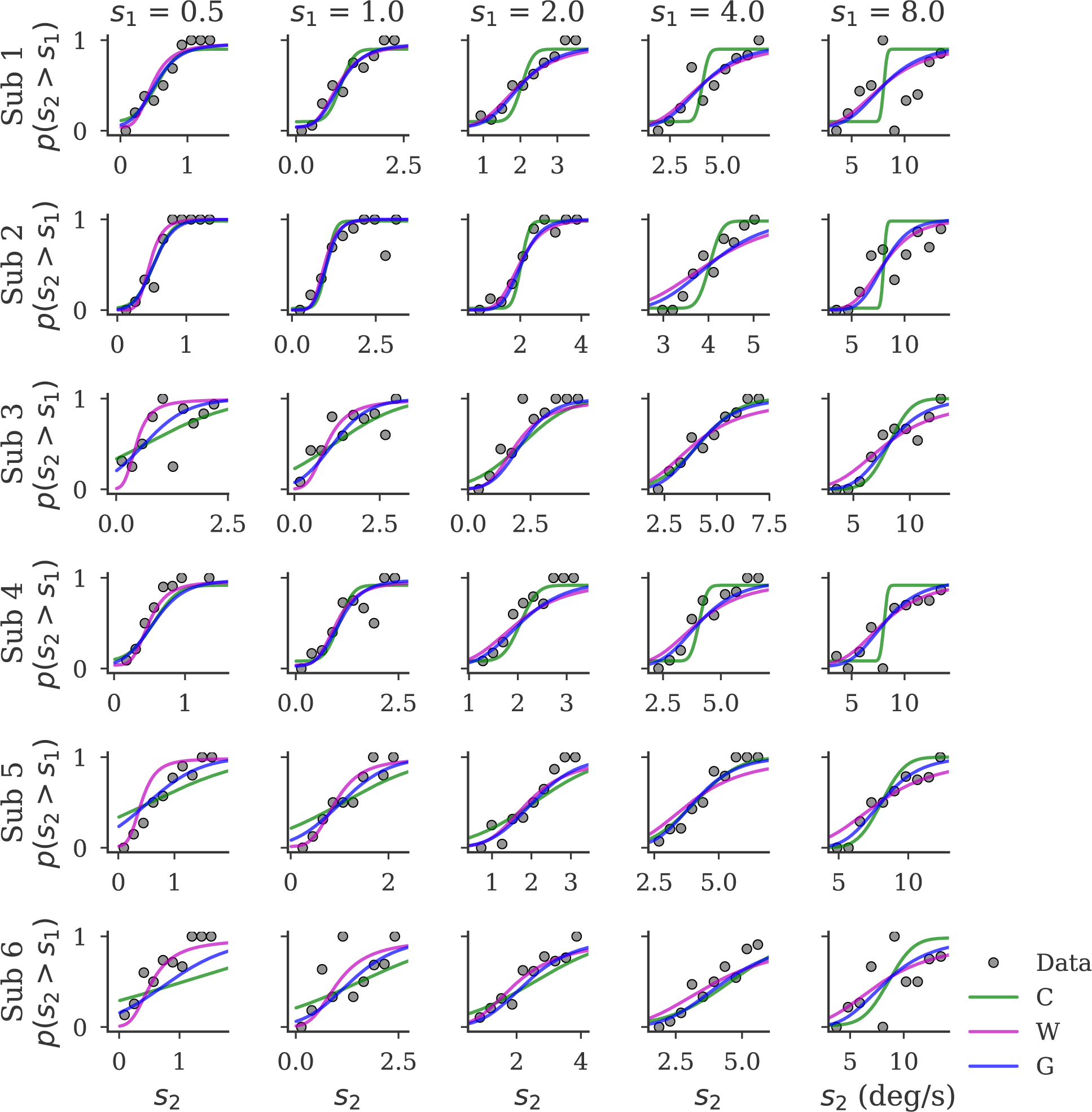
Data and model predictions for psychometric curves measured in the psi session. Each row indicates the psychometric curves of a particular subject, each column indicates the reference value (*s*_1_) for this psychometric curve. Grey dots indicate proportion of trials where observers report *s*_2_ *> s*_1_, proportions were obtained by binning responses in 10 bins from the minimum to maximum probe value (*s*_2_) for this subject and reference (*s*_1_) value. Curves indicate the predicted proportion from each of the models. Note, the parameters used for the predictions were obtained from fitting only to stimuli selected using our algorithm and thus were not fit to the data shown.

Another important property of adaptive algorithms is that they do not sample uniformly across the entire stimulus space. Instead, the stimuli selected are those that are most informative to compare the models. In order to visualize which stimuli these are in this experiment we plotted the stimuli selected using the adaptive method for a representative subject (see Figure 6). The adaptive sampling method alternates between high and low speeds for the reference and probe stimuli. This sampling strategy is sensible as the noise models make distinct predictions for high and low speeds and thus sampling at high and low speeds allows for effective dissociation of the models.

**Figure 6.**
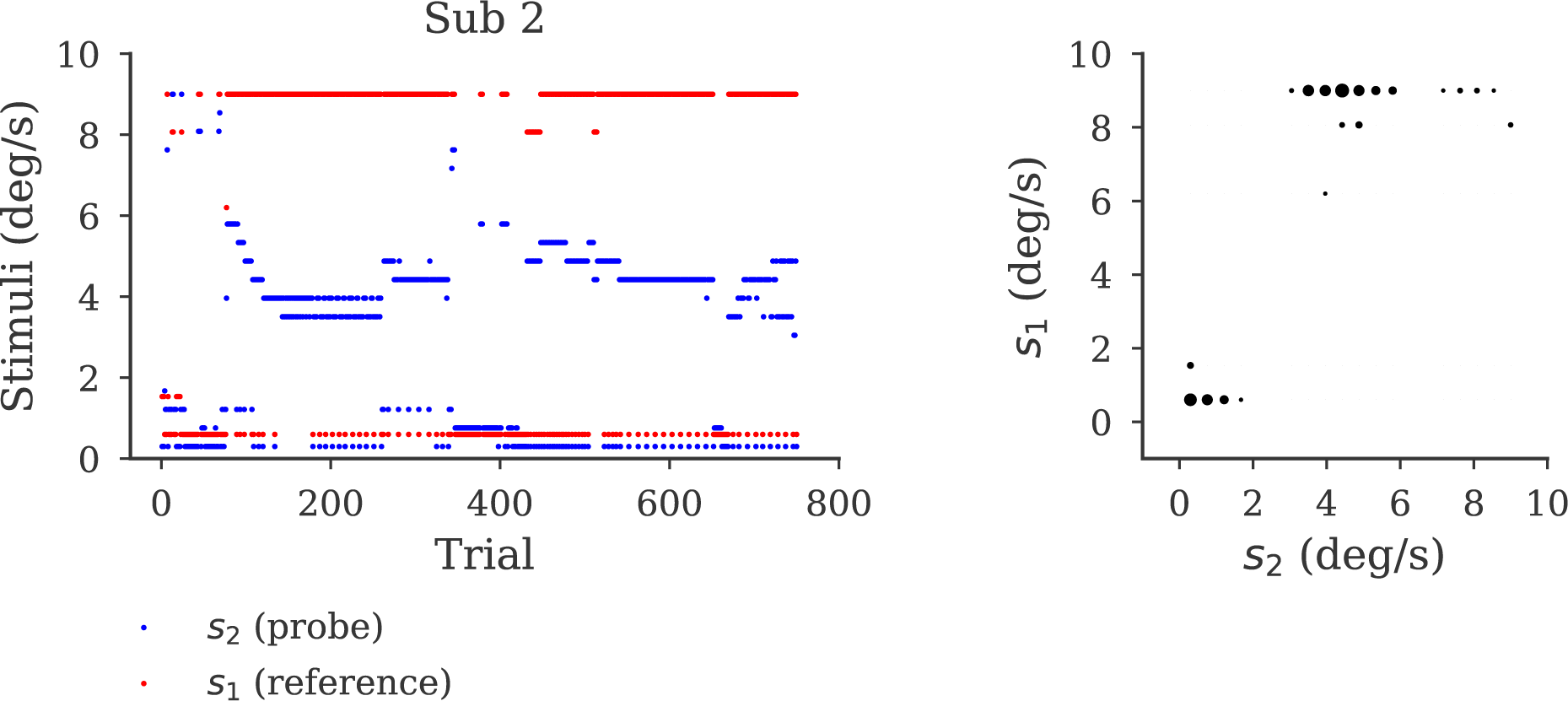
Stimuli adaptively selected for subject 2. The left plot shows the probe (blue dots) and reference (red dots) selected on each trial in experiment 1. The right plot shows a scatter plot of the combination of probe and reference. Radius of the data points is proportional to 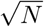 where *N* number of times this combination was selected.

## Experiment 2: Target selection

### Introduction

The previous section illustrates the use of our algorithm as a method to dissociate different sensory noise models. However, this is only one example comparison. To ensure our algorithm is broadly applicable, it is important to validate it in multiple settings. Here, as an additional application, we consider comparing models of saccadic target selection during self-motion (Rincon-Gonzalez et al., 2016), a study recently performed in our lab. This example allows us to investigate how much benefit our algorithm provides when the models being compared are highly non-linear and the signal-to-noise ratio in the data is low. In this experiment, subjects were passively translated from left to right in a sinusoidal motion profile and at 8 pre-defined phases of the oscillation two targets were presented. The subjects were instructed to make a saccade to one of the two targets, which were presented asynchronously with a particular stimulus onset asynchrony (SOA). This produces a single psychometric curve of subject’s choice as a function of SOA for each phase. This curve can then be used to determine the SOA at which the probability of selecting each target is equal, referred to as the balanced time delay (BTD). The experiment showed that, on the group level, BTD changes sinusoidally as a function of the motion phase suggesting that subject’s target selection behavior, and thus preference, is influenced by current body motion. However, the amplitude of the modulation was small and the signal-to-noise ratio was low, which made comparing a sinusoidal modulation to alternative models difficult at the individual subject level. Our algorithm may provide a solution to this difficulty, as adaptive sampling selects the most informative stimuli to dissociate the selected models.

Here, we first reanalyze data from this experiment and show that the data of approximately half of the subjects are best described by a sinusoidal modulation rather than a constant choice bias. In other subjects the results of the model comparison are inconclusive. We next demonstrate with simulations that using our algorithm for stimulus selection would have improved model comparison accuracy. This suggests our algorithm is also useful to help dissociate models in circumstances where the signal-to-noise ratio is limited.

## Methods

### Models

In order to test whether self-motion has any effect on psychophysical choice behavior we consider two models of choice behavior, a constant bias model and a sinusoidal bias model (Bakker et al., 2017). We model choice behavior as:

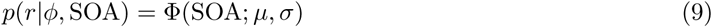

in which *r* is the subject’s response, *φ* is the phase at which the targets are presented, Φ is a cumulative Gaussian with mean *µ* and standard deviation *σ* evaluated at the *SOA*. For the constant model, *µ* is a fixed value across phases: *µ* = *α*. In this model choices are independent of the phase of the motion. The sinusoidal model entails *µ* changes sinusoidally as a function of phase, and is thus written *µ* = *α* + *β* sin(*φ* + *φ_o_*), in which *α*, *β* and *φ_o_* are free parameters representing a subject’s fixed bias, amplitude of the modulation and phase offset, respectively. Regardless of the model, we can parameterize the subject response probability using *θ* = [*α, β, φ_o_, σ*].

### Reanalysis

In order to test whether the individual subject’s choice behavior is modulated sinusoidally and to obtain reasonable parameters to utilize in our simulations we reanalyzed the data of 17 subjects from Rincon-Gonzalez et al. (2016). We fit both the sinusoidal and constant bias models to each subject’s choice data. We assumed the responses are independent across trials. The response probability on each trial can be computed using equation 9. The log-likelihood of a subjects’ data set is then,

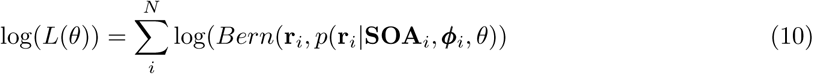

in which *i* is the trial index, *N* is the number of trials, **r** is a vector of subject responses, **SOA** is a vector of the SOA’s the subject was presented, ***φ*** is a vector containing the phase the targets were presented at and *Bern* stands for a Bernoulli distribution.

Parameter estimates 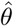 were then obtained using equation 7. As before this optimization was done numerically using the L-BFGS-B algorithm (Byrd et al., 1995). The parameter bounds were set to those in Table 4. To ensure a global minimum was found we used 100 random initializations and selected the parameter set with the highest log-likelihood. The initial values were obtained by drawing each parameter value from a continuous uniform distribution with the same bound as those in Table 4.

**Table 4.** Parameter grids used for simulation experiment 2.

| Variable | Lower bound grid | Upper bound grid | Number of steps |
| --- | --- | --- | --- |
| $\phi$ (rad) | 0 | 5.5 | 8 |
| $SOA$ (ms) | -250 | 250 | 25 |
| $\alpha$ (ms) | -70 | 70 | 15 |
| $\beta$ (ms) | 0 | 60 | 15 |
| $\phi_o$ (rad) | -3 | 3 | 15 |
| $\sigma$ (ms) | 50 | 190 | 15 |

In order to validate the results of the grid-based model comparison we also computed the Akaike Information Criterion (AIC) for each of the models using equation 8. As an additional analysis we fit a cumulative Gaussian (see equation 5) to the data from each phase (using the same bounds as for the constant model and *λ* set to 0) to provide us with a semi-parametric estimate of BTD for each phase.

### Simulation experiment

In order to investigate whether using our adaptive algorithm could help to dissociate these different models of target selection, we performed a simulation experiment. The required grids are specified in Table 4. As priors we used a uniform discrete distribution for each parameter and a uniform distribution over the two models.

We first generated 2000 possible parameter combinations. Parameters were drawn independently from a continuous uniform distribution with the same upper and lower bound as those specified in Table 4. Next, in order to assess how well we can infer the correct generative model for each parameter combination we simulated a synthetic subject performing 1000 trials for each generative model and parameter combination. Note, the constant model is independent of *β* and *φ*_0_ and thus they were removed from the parameter set when simulating this model. The stimuli for these trials were selected either randomly from the stimulus grid shown in Table 4 or using our adaptive algorithm. This led to a total of 8000 simulated datasets. Additional simulations were performed based on a truncated Gaussian parameter distribution, reflecting the estimated behavioral parameter range (see Supplementary material).

## Results

The AIC scores and parameter estimates for both models are shown in Table 5. In order to interpret the AIC scores it is useful to note that an AIC difference of over 4 is considered positive evidence towards the model with the lower score (Burnham et al., 2002). This suggests the model comparison in 8 of the subjects is ambiguous (AIC difference under 4), no subjects are best described by the constant bias model and 9 subjects are best described the sinusoidal bias model. Interestingly, it can be seen that even in the ambiguous cases the amplitude parameter *β* is not at zero. This implies the modulation of BTD is sinusoidal but the effect on the log-likelihood is insufficient to overcome the penalization for the additional parameters. This is also supported by the model predictions shown in Figure 7, illustrating that the sinusoidal model is a closer fit than the constant model to the independent estimate of BTD for each phase.

**Table 5.** Best fit parameters and AIC differences (ΔAIC, sinusoidal - other model) for each model for all subjects.

| Subject | Model | $\alpha$ (ms) | $\beta$ (ms) | $\phi_o$ (rad) | $\sigma$ (ms) | $\Delta\text{AIC}$ |
| --- | --- | --- | --- | --- | --- | --- |
| 1 | Sinusoidal Bias | 67.478 | 15.895 | -0.002 | 74.505 | 0.000 |
| 1 | Constant Bias | 67.467 | N/A | N/A | 76.780 | -8.469 |
| 2 | Sinusoidal Bias | -69.792 | 19.323 | -1.653 | 158.262 | 0.000 |
| 2 | Constant Bias | -69.589 | N/A | N/A | 160.475 | -2.810 |
| 3 | Sinusoidal Bias | -14.556 | 8.692 | 1.436 | 65.669 | 0.000 |
| 3 | Constant Bias | -14.492 | N/A | N/A | 66.621 | -1.480 |
| 4 | Sinusoidal Bias | 42.007 | 47.468 | -1.161 | 176.735 | 0.000 |
| 4 | Constant Bias | 42.793 | N/A | N/A | 190.810 | -23.854 |
| 5 | Sinusoidal Bias | -25.052 | 1.283 | 0.607 | 61.613 | 0.000 |
| 5 | Constant Bias | -25.050 | N/A | N/A | 61.623 | 3.871 |
| 6 | Sinusoidal Bias | -54.271 | 15.12 | -0.481 | 65.382 | 0.000 |
| 6 | Constant Bias | -54.346 | N/A | N/A | 67.457 | -11.753 |
| 7 | Sinusoidal Bias | 21.550 | 24.817 | -1.21 | 68.076 | 0.000 |
| 7 | Constant Bias | 21.652 | N/A | N/A | 73.418 | -30.514 |
| 8 | Sinusoidal Bias | 12.830 | 14.322 | 1.012 | 95.736 | 0.000 |
| 8 | Constant Bias | 12.836 | N/A | N/A | 97.677 | -3.968 |
| 9 | Sinusoidal Bias | 0.216 | 16.564 | -0.529 | 97.097 | 0.000 |
| 9 | Constant Bias | 0.318 | N/A | N/A | 99.491 | -4.859 |
| 10 | Sinusoidal Bias | 3.735 | 15.64 | -0.956 | 127.027 | 0.000 |
| 10 | Constant Bias | 3.783 | N/A | N/A | 129.383 | -1.729 |
| 11 | Sinusoidal Bias | 60.449 | 13.294 | -0.295 | 116.987 | 0.000 |
| 11 | Constant Bias | 60.467 | N/A | N/A | 118.368 | -0.652 |
| 12 | Sinusoidal Bias | 7.974 | 16.892 | 0.394 | 117.834 | 0.000 |
| 12 | Constant Bias | 8.233 | N/A | N/A | 119.763 | -3.079 |
| 13 | Sinusoidal Bias | -1.041 | 19.071 | -0.359 | 74.627 | 0.000 |
| 13 | Constant Bias | -1.056 | N/A | N/A | 77.387 | -16.147 |
| 14 | Sinusoidal Bias | -27.310 | 17.567 | -0.847 | 151.860 | 0.000 |
| 14 | Constant Bias | -27.203 | N/A | N/A | 154.390 | -0.969 |
| 15 | Sinusoidal Bias | -13.869 | 10.955 | -0.759 | 68.580 | 0.000 |
| 15 | Constant Bias | -13.752 | N/A | N/A | 69.499 | -5.662 |
| 16 | Sinusoidal Bias | -4.454 | 48.497 | 1.085 | 122.325 | 0.000 |
| 16 | Constant Bias | -4.918 | N/A | N/A | 141.447 | -56.525 |
| 17 | Sinusoidal Bias | 39.354 | 14.658 | 0.227 | 68.175 | 0.000 |
| 17 | Constant Bias | 39.470 | N/A | N/A | 70.346 | -12.939 |

**Figure 7.**
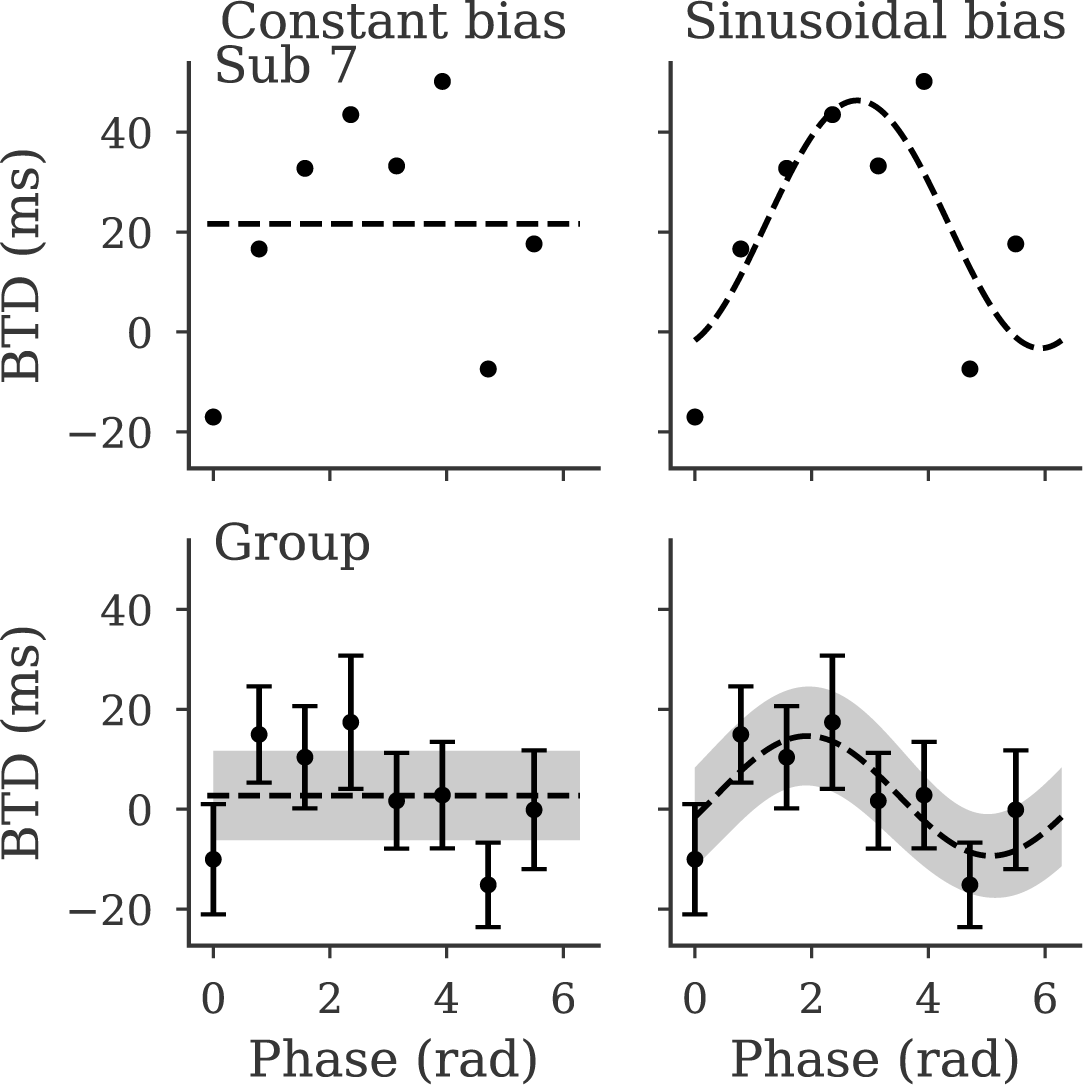
Sinusoidal and constant model predictions for an example subject and across subjects. For the group the dashed line indicates the mean predicted BTD across subjects, for the example subject it indicates the predicted BTD. The shaded regions indicates ± 1 *SEM* across subjects. Data points are the BTD obtained by fitting a psychometric curve to each phase. The error bars indicate ± 1 *SEM* across subjects.

In order to explore if our algorithm can facilitate model comparison, we plotted the average model probabilities across trials for both models and sampling methods used in our simulation experiment (see Figure 8). The model probabilities trend to 1 along the diagonal, indicating both adaptive and random sampling converge towards the correct model. As before, the probabilities are higher for the adaptive sampling method compared to random sampling suggesting that our algorithm increases the strength of evidence towards the correct model. The magnitude of this increase is lower than observed in simulation experiment 1.

**Figure 8.**
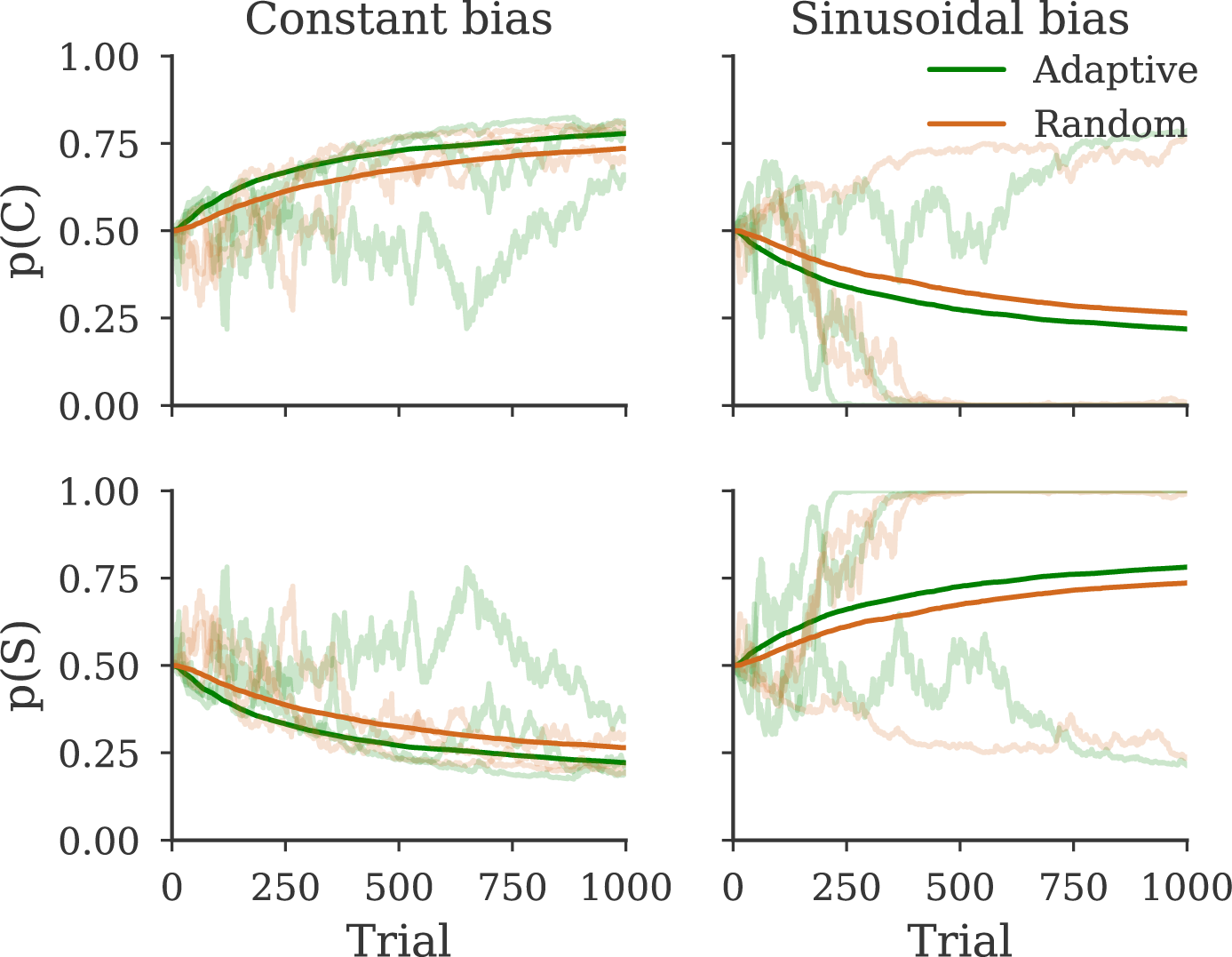
Evolution of model probabilities over trials for different generative models and algorithms. Columns indicate the model used to generate the data, rows indicate the probability of each model. The dark lines indicate the mean probability averaged over simulations, light lines indicate example simulations. Green coloring indicates stimuli were selected adaptively, orange coloring indicates stimuli were selected at random from the same stimulus grid.

We also quantified how each sampling method affects the conclusions drawn by computing the Bayes factor of the generative model against the other model. These Bayes factors are plotted in Figure 9. Interestingly, if stimuli are selected randomly and the correct model is sinusoidal we only conclude in favor of it in 60% of the simulations. This matches with the mixed results from the reanalysis. Adaptive sampling increases the proportion of simulations in which we find strong evidence in favor of the correct model. For the sinusoidal model, we obtained a benefit of about 15%, which is a smaller benefit than observed in the noise model simulation.

**Figure 9.**
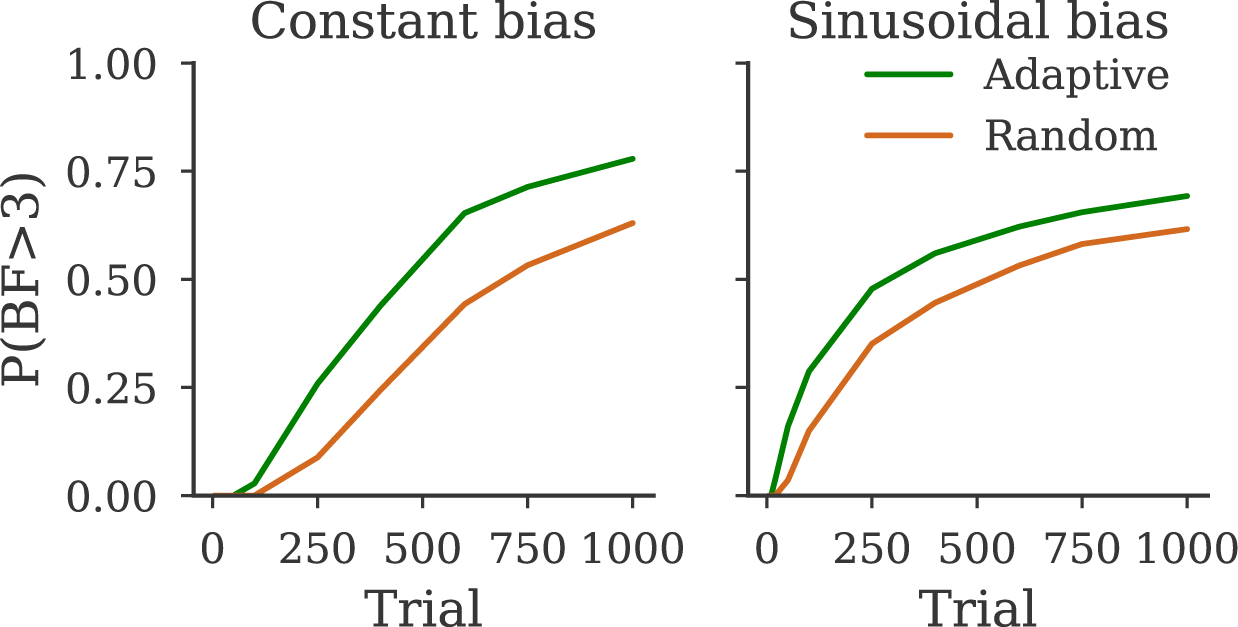
Proportion of simulations where the Bayes factors with respect to the generative model is over 3. Each column indicates the model used to generate the data.

In order to explore why the models cannot be strongly dissociated in each simulation, we plotted the probability of the correct model as a function of *σ* and *β* (see Figure 10). If the generative model is the constant bias model, both adaptive and random sampling method lead to model probabilities favoring the correct model but the adaptive method produces only slightly higher probabilities. Adaptive sampling leads to the probability of the correct model being slightly higher (indicated by a more yellow hue), which leads to a larger proportion of Bayes factors being over 3. By contrast, when the generative model is the sinusoidal model, the model probabilities range from strongly in favor of the sinusoidal model to strongly in favor of the constant model for both sampling methods. This is understandable because the smaller the amplitude of the sinusoid, the closer the sinusoidal model becomes to the constant model and thus penalizing for the additional parameters leads to favoring the simpler constant model. Interestingly, the shift in model probabilities from sinusoidal to constant is dependent on the variability of a subjects decisions; the smaller *σ* is, the lower *β* can be, while still inferring in favor of the sinusoidal model.

**Figure 10.**
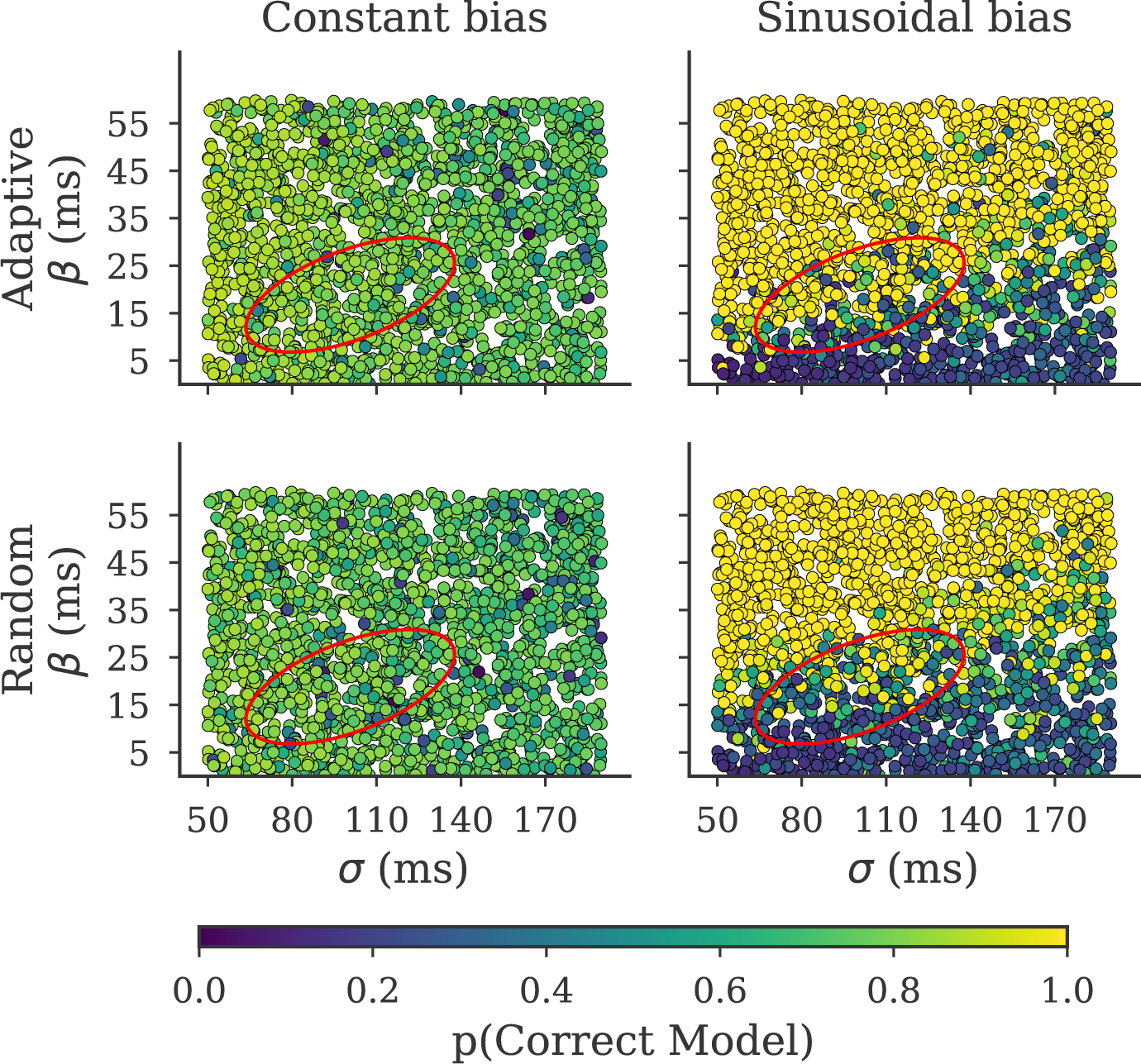
Probability of the generative model as a function of parameter values for different generative models and algorithms. Columns indicate the model used to generate the data, rows indicate the sampling method used to determine stimuli. Each point indicates the probability of the correct model as a function of amplitude *β* and standard deviation of a subject’s choices *σ*. Note, constant bias model is independent of amplitude *β*, thus model probabilities do not change systematically as a function of *β*. The *β* value plotted refers to the value used in the sinusoidal model. The red ellipse indicates the mean ± one standard deviation of the subject’s parameters obtained from the sinusoidal model (see Table 5).

To determine why adaptive sampling improves the chance of inferring in favor of the correct generative model, Figure 11 illustrates the phase and SOA selected using the adaptive algorithm for an example simulation. In the initial trials, the algorithm samples broadly over the phase and SOA, but then converges to a few combinations of SOA and phase. Specifically, adaptive sampling selects the phases where the BTD is maximal or minimal and SOA values close to the current *α* estimate.

**Figure 11.**
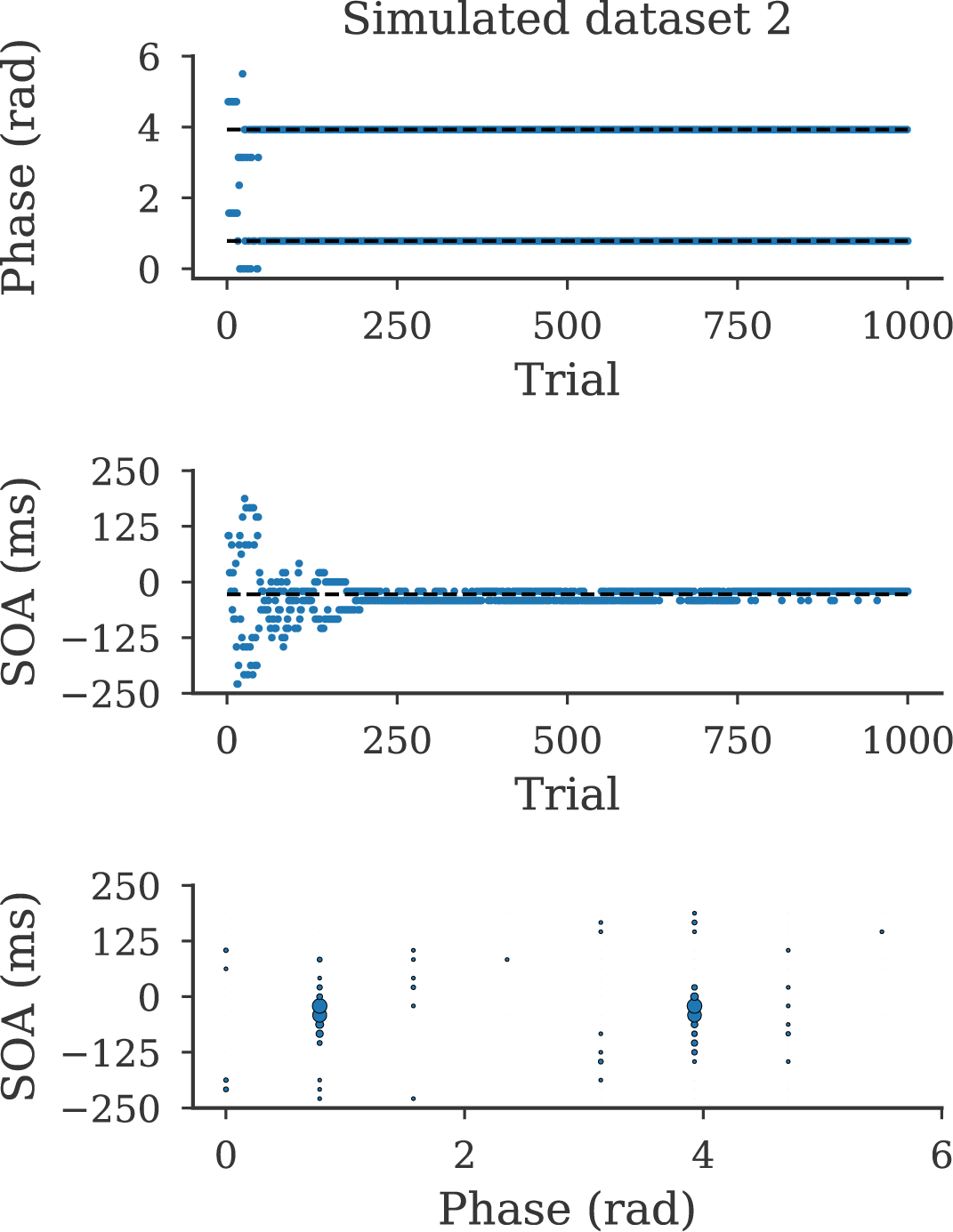
Stimuli selected adaptively for an example simulation. The upper two plots indicate the phase and SOA sampled across trials. In both plots the blue dots indicate the sampled stimuli for a particular trial. For the phase plot the dashed lines indicate the phases (from our stimulus set) for which the BTD is maximal or minimal. For the SOA plot the dashed line indicates the baseline BTD (the BTD independent of phase modulations). The lower plot indicates a scatter plot of the combination of phase and SOA. The radius of the data point is proportional to 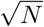 where *N* is the number of times this combination was selected.

## Discussion

Using a series of simulations in which the correct generative model is known, we show that selecting stimuli adaptively increases the probability of inferring the correct generative model. We further show this increase affects the conclusions an experimenter could draw. When stimuli are selected adaptively an experimenter is more likely to conclude strongly in favor of the correct generative model and it requires fewer trials to reach this conclusion. For example, in Figure 2 when the generative model is the generalized model, our adaptive algorithm yields in only 250 trials strong evidence towards the correct model in 60% of the simulations. By contrast almost none of the simulations using random sampling showed strong evidence.

We illustrate this model comparison benefit in two distinct settings, firstly, dissociating different sensory noise models and secondly, dissociating models of target selection. As an additional step towards practical application, we also used our algorithm to test between sensory noise models of human speed perception. We found that selecting stimuli adaptively increases the strength of evidence towards the model previously proposed (Stocker & Simoncelli, 2004, 2006).

Our findings match with previous work in cognitive science which illustrate that models of memory retention can be better dissociated by selecting stimuli adaptively (Cavagnaro et al., 2010, 2011). We also illustrate that the magnitude of improvement provided by adaptive sampling is highly specific to the models being compared. Specifically, we found a dramatic improvement in dissociating sensory noise models but only a small improvement in dissociating models of saccadic target selection.

Being able to compare models in an efficient manner encourages comparison of different models which may otherwise not be compared. For example, in many cases the sensory noise model is a single component of a more complex model (Acerbi et al., 2012, 2017; Jazayeri & Shadlen, 2010). In the aforementioned work, the possibility of different sensory noise models is dealt with through model comparison. However, incorporating multiple sensory noise models adds an additional degree of freedom, to the space of possible models, which can introduce difficulties in model comparison (Acerbi, Ma, & Vijayakumar, 2014). Specifically, multiple models with different components (for instance, sensory noise, priors, loss functions) can fit the same data equally well which makes inferring the correct components difficult (Acerbi et al., 2014). This study also indicated a possible solution to this problem; fixing certain model components and parameters based on previous work or additional experiments. As such an experimenter could perform an additional experiment to test the sensory noise model (and also obtain parameter estimates) for each subject, which could then be fixed in the model comparison. Our algorithm presents an efficient way to test between the noise models in a small number of trials and therefore could be used as a method for efficient model selection.

Although we illustrate, in two distinct practical examples, the benefits of using our algorithm, there are limitations to our approach. One major limitation is the grid-based approach we use in our algorithm. While this approach is reasonable for the relatively simple models we tested here. It is unfeasible for more complex models (models with either more parameters or more stimuli dimensions). This is because if we use the same sized grid for each parameter the number of points increases exponentially with the number of parameter dimensions or stimuli dimensions (DiMattina, 2015). For more complex models, these grids could exceed the RAM memory available in certain computers, preventing our algorithm from being applicable. In addition, more complex models will require more time to compute the optimal stimulus. For example, it takes approximately 100 ms with our current models, the additional time increase may render the current implementation unfeasible for more complex models. Fortunately, there are a number of different approaches which can compensate for these problem. One method is to use an adaptive approach to selecting the number of grid points and their positions (Kim, Pitt, Lu, Steyvers, & Myung, 2014; Pflüger, Peherstorfer, & Bungartz, 2010). The notion is that the contribution of each point in the parameter space is not equal and thus more points should be used for more informative regions of the parameter space. This approach, previously suggested in the context of parameter estimation (DiMattina, 2015) could allow our algorithm to scale to higher dimensional models, or to more than three models. Another alternative solution is to use an analytic approximation to the parameter posterior, for example by using a Laplace approximation (DiMattina, 2015) or by a sum-of-Gaussians (DiMattina & Zhang, 2011) and compute the optimal stimuli based on the approximated posterior. With such an approximation it is only necessary to maintain the parameters for the approximation rather than large grids. Again allowing our algorithm to scale up to higher dimensions and more models. However, this comes at the computational cost of having to refit each of these approximations to every model on each trial. As the time required to evaluate the likelihood typically increases approximately linearly with the number of datapoints, this means the time required to refit these approximation increases with the duration of the experiment (DiMattina, 2015). Additionally, if the shapes of the posteriors are a poor match to these approximations (for example, highly skewed distributions are poorly approximated using a Laplace approximation) then this approach may perform poorly compared to grid approximations which present a non-parametric method of representing the posterior (DiMattina, 2015). Given that these different approaches have distinct costs and benefits, it is important to quantitatively test them to see how each performs in terms of accuracy, computation time and memory usage. A detailed comparison of this type has been performed in terms of adaptive stimulus selection for parameter estimation (DiMattina, 2015), but to our knowledge, no such analysis has been performed for model comparison. An important avenue for further work would be to explicitly compare our algorithm to other existing algorithms (DiMattina, 2016; Cavagnaro et al., 2010) to identify the relative costs and benefits of each approach.

In addition to the practical limitations of our approach, it is important to consider the theoretical implications of using adaptive sampling on model comparison. For example, adaptive sampling could significantly change the distribution of stimuli presented to the subjects (see Figure 6) and therefore could violate assumptions used in certain model comparisons. For example, it is assumed the subject’s underlying model is independent of the stimuli presented and typically Bayesian observer models assume that the subject’s priors matches the stimulus distribution (Keshvari et al., 2012, 2013). The adaptive approach may cause violations of these assumptions. To illustrate this, consider the change detection experiments referenced above (Keshvari et al., 2012, 2013). In these experiments subjects are first shown a number of oriented ellipses which the subject has to memorize. Subsequently the ellipses are displayed again either with the same orientation or a changed orientation and the observer must report whether a change is perceived or not. All the models compared for this task assumed the subject used a circular uniform prior over the size of the change (the same used to generate the stimuli). If we were to generate the change magnitude adaptively instead, this could create a non-uniform distribution. Presenting a non-uniform distribution of change magnitude may cause subjects to alter their response strategy. For example, if a subject is only being presented trials with large changes he/she may shift from encoding the stimuli precisely to a more coarse encoding of the stimuli as precise encoding is no longer needed for the task. This biased distribution could also create a mismatch between the assumed (uniform circular) prior in the model and the actual experiment, which could cause biases in model comparison.

Although these issues may seem severe, the risk can be mitigated. Our suggestion is to not rely only on adaptive techniques as definitive evidence towards a model. It is important that multiple experiments and sampling methods support the same model. In some cases discrepancies may be found between sampling methods (e.g. in our noise model comparison experiment). In these cases it is important to perform simulations to see if these results are to be expected (see Supplemental material for the simulation we performed) or if the adaptive technique could be biasing the comparison.

A final theoretical point is that our algorithm assumes the true model used by the subject is part of the included set of models being considered (an assumption in all parametric model comparisons). If the true model is not part of this set then the stimuli are not optimized to find evidence for this model. Obviously, in real subjects, it is impossible to know what the ‘true’ model is, rather we are searching for realistic models that best explain the subject data. It is important to be aware that when using any adaptive approach the stimuli are only optimized for dissociating the assumed model set.

An additional area for further work is the importance of priors in dissociating models. For simplicity, we used uniform priors for both models and parameters. However, this neglects prior information which may reduce the number of trials necessary to estimate which model is the best. How should we determine these priors? Within statistics itself there is little consensus on how this should be done, ranging from the prior being a subjective choice of the experimenter (de Finetti, 2017) to the prior being objectively estimated from data (Jaynes, 2003). Recent work has embraced the latter approach and used hierarchical Bayesian modeling to estimate the prior based on previous subjects (Kim et al., 2014). For example, this approach has been successful in determining parameter priors to use for observer’s contrast sensitivity functions, both in simulations and in actual experiments (Gu et al., 2016; Kim et al., 2014). A similar approach could be taken for estimating both parameter and model priors by creating a hierarchical model which incorporates the different models to be compared and fitting this to data from previous subjects. An important step for further work would be to formalize this generalization and investigate how these priors affect model inference.

## Acknowledgments

This work was supported by the European Research Council grant EU-ERC-283567 (to WPM) and the Netherlands Organization of Scientific Research grants NWO-VICI: 453-11-001 (to WPM and JRHC) and NWO-VENI: 451-10-017 (to LPJS). We would like to thank the two anonymous reviewers for their helpful comments, including the change detection example.

## Appendix A

In order to model a subject’s 2-afc behavior as a function of different sensory noise models we assume a subject receives two sensory measurements *x*_1_ and *x*_2_, one for the reference and one for the probe. We model these as normally distributed random variables, with a mean centered on the true reference and probe values and a variance which is a function of the underlying sensory noise model. As such we can write *x*_1_ and *x*_2_ as 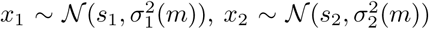. We assume an observer responds 1 if *x*_2_ *> x*_1_ and 0 otherwise. In order to derive the distribution of an observer’s response it is useful to note this is equivalent to *x*_2_ − *x*_1_ *>* 0. As *x*_2_ and *x*_1_ are normal distributed random variables, subtracting them produces another normally distributed variable ∆_*x*_. Therefore the subject’s response probability can be written as:

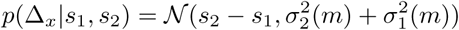

The likelihood of an observer responding 1 is obtained by integrating over positive values of ∆_*x*_,

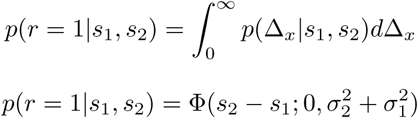

Because the responses are mutually exclusive, it follows that the likelihood of a subject responding 0 is,

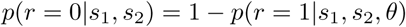

in which Φ is the cumulative normal distribution, evaluated at point *s*_2_ − *s*_1_, with a mean of 0 and variance 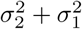. This entails that a subject’s 2-afc behavior is unbiased and also that subjects do not lapse during the experiment. To make the model more realistic, we augment it with a small bias term *α* to account for small deviations from unbiased behavior and a lapse term *λ* to account for lapses in the task. Therefore the final response probability can be written,

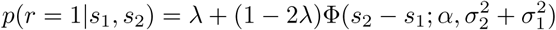

It is important to note that subjects do not estimate the underlying speed (using Bayes rule) as they are using sensory observations rather than posterior estimates. This was done for two reasons. Firstly, there is not a consensus on how additional information is incorporated in speed perception; some models propose that observers incorporate assumptions about motion dynamics to create priors (Kwon, Tadin, & Knill, 2015), others propose statistics of natural stimuli are used to form priors (Stocker & Simoncelli, 2004, 2006). Secondly, unless a uniform prior (across the real line) or conjugate prior is used, computing the response probability in closed form is difficult (recent work has analytically derived the effect of Gaussian priors in 2-afc tasks (Acuna, Berniker, Fernandes, & Kording, 2015)).

Because our main focus is the sensory noise model, not the incorporation of priors, our experiment was designed such that the influence of priors should be negligible and hence our derived response probability should be a reasonable approximation. Specifically, it has been shown that the bias in speed estimation decreases when stimuli are close to fixation (Kwon et al., 2015) and biases decrease when contrast is high (Stocker & Simoncelli, 2004, 2006). Our stimuli were centered relatively close to fixation (6 deg eccentricity) compared to other speed perception experiments (Kwon et al., 2015) and also had a much higher contrast than is typical (Stocker & Simoncelli, 2004, 2006). This means most of subject behavior should be governed by the sensory noise and not the prior.

